# Short periods of bipolar anodal TDCS induce no instantaneous dose-dependent increase in cerebral blood flow in the targeted human motor cortex

**DOI:** 10.1101/2022.01.10.475608

**Authors:** Marie Louise Liu, Anke Ninija Karabanov, Marjolein Piek, Esben Thade Petersen, Axel Thielscher, Hartwig Roman Siebner

## Abstract

**Background:** Anodal transcranial direct current stimulation (aTDCS) of primary motor hand area (M1-HAND) can enhance corticomotor excitability. Yet, it is still unknown which current intensity produces the strongest effect on regional neural activity. Magnetic resonance imaging (MRI) combined with pseudo-continuous Arterial Spin Labeling (pc-ASL MRI) can map regional cortical blood flow (rCBF) and may thus be useful to probe the relationship between current intensity and neural response at the individual level.

**Objective:** Here we employed pc-ASL MRI to map acute rCBF changes during short-duration aTDCS of left M1-HAND. Using the rCBF response as a proxy for regional neuronal activity, we investigated if short-duration aTDCS produces an instantaneous dose-dependent rCBF increase in the targeted M1-HAND that may be useful for individual dosing.

**Methods:** Nine healthy right-handed participants received 30 seconds of aTDCS at 0.5, 1.0, 1.5, and 2.0 mA with the anode placed over left M1-HAND and cathode over the right supraorbital region. Concurrent pc-ASL MRI at 3 T probed TDCS-related rCBF changes in the targeted M1-HAND. Movement-induced rCBF changes were also assessed.

**Results:** Apart from a subtle increase in rCBF at 0.5 mA, short-duration aTDCS did not modulate rCBF in the M1-HAND relative to no-stimulation periods. None of the participants showed a dose-dependent increase in rCBF during aTDCS, even after accounting for individual differences in TDCS-induced electrical field strength. In contrast, finger movements led to robust activation of left M1-HAND before and after aTDCS.

**Conclusion:** Short-duration bipolar aTDCS does not produce instantaneous dose-dependent rCBF increases in the targeted M1-HAND at conventional intensity ranges.

Therefore, the regional hemodynamic response profile to short-duration aTDCS may not be suited to inform individual dosing of TDCS intensity.

**Highlights:**

- Arterial spin labeling (ASL) MRI probed regional cerebral blood flow (rCBF) during anodal TDCS.
- Short-duration anodal TDCS was applied to left motor hand area (M1-HAND) at 0.5, 1.0, 1.5 and 2.0 mA.
- Anodal TDCS produced no instantaneous dose-dependent rCBF increase in left M1-HAND.
- Apart from a subtle increase at 0.5 mA, rCBF was not modified in M1-HAND during anodal TDCS.

## Introduction

Transcranial Direct Current Stimulation (TDCS) is widely used to modulate cortical function in cognitive neuroscience and in therapeutic settings [1–8]. The underlying mechanism of action still remains to be fully clarified. Neurophysiological studies suggest that TDCS modulates the intrinsic activity of cortical neurons by inducing shifts in the neuronal membrane potential, presumably by polarizing axon terminals [9,10].

In therapeutic settings, the majority of TDCS studies have yielded varying results and have failed to provide clear evidence for efficacy [11]. High inter-individual variability has also been observed in neurophysiological studies which used single-pulse Transcranial Magnetic Stimulation (TMS) and recordings of the motor evoked potential (MEP) to probe the aftereffects of TDCS on corticospinal excitability [12]. Using a bipolar montage with one skin electrode placed on the scalp over the primary motor hand area (M1-HAND) and the other electrode placed on the contralateral supraorbital region, TDCS produced polarity-specific bi-directional effects on corticomotor excitability [13–15]. The consistency of the after-effects has been challenged by several recent studies, which found substantial inter-individual variability of the TDCS aftereffects without a clear excitability shift at the group level [16–18].

The large inter-individual variability may partially be caused by the way current intensity of TDCS is determined. Differences in head and brain anatomy significantly influence strength and spatial distribution of the induced electrical field (E-field) [19,20]. Adjusting the stimulation intensity using field simulations informed by individual structural MRI can account for some variance at the single-person level [21,22] but we still lack personalized “dosing methods” that allow to adjust the current intensity of TDCS based on the individual cortical response profile. Personalized dosing requires reliable brain mapping approaches that can reliably delineate the dose-response relationship between the TDCS current intensity and the TDCS induced change in regional cortical activity at the single-person level. This information might help to personalize the current intensity of TDCS in a way that the induced E-fields optimally engages the targeted cortical area.

One way to study how TDCS modulates regional neuronal activity is to map the regional neurovascular response to TDCS with functional magnetic resonance imaging (MRI). Functional MRI (fMRI) exploits the blood oxygen level dependent (BOLD) contrast, while Arterial Spin Labelling (ASL) uses magnetically labeled water in the arterial blood as a tracer of regional cerebral blood flow (rCBF) [23,24]. Most BOLD fMRI studies have examined after-effects of longer periods of TDCS [25–29]. However, immediate changes in neural activity induced during short periods of TDCS reflect more directly how TDCS impacts on neuronal activity in the cortical target region. This notion is supported by TMS studies which showed that short periods of bipolar anodal TDCS (4-120 seconds) targeting M1-HAND induce an immediate rise in cortical excitability, indicated by an increase in MEP amplitude, without sustained after-effects [13,30]. Animal studies further support the immediate increase in cerebral perfusion during transcranial electric stimulation, as dose-dependent blood flow changes rapidly follow the initiation and termination of short-duration (6-30 seconds) TACS [31].

Despite of the evidence for immediate neurovascular effects of TDCS, fMRI studies focusing on immediate TDCS effects are sparse [32–34]. Perfusion studies measuring sustained effects of TDCS with longer stimulation periods indicate a general increase of regional cerebral blood flow (rCBF) during bipolar TDCS [29,35,36] and recent ASL-fMRI studies show dose-related increases in rCBF in the cortex underneath the precentral electrode during and after anodal TDCS at conventional intensities (0.5 – 2.0 mA) [37] and at higher intensities up to 4mA [38].

Prompted by these studies, we performed concurrent ASL-MRI during bipolar anodal TDCS (aTDCS) of left M1-HAND to test whether short periods of aTDCS trigger dose-dependent changes in rCBF in the targeted M1-HAND. We were particularly interested to examine whether concurrent ASL-MRI reliably reveal a dose-response profile at the single-subject level. We used the classic bipolar montage that has been used to induce changes in corticospinal excitability in the M1-HAND and administered 30-seconds epochs of bipolar aTDCS at stimulation intensities from 0.5 – 2.0 mA [39]. We hypothesized that the rCBF increase induced by short-duration aTDCS would linearily and positively scale with TDCS current intensity, as it has been shown for TDCS protocols applied over a longer stimulation period [37].

## Material and methods

### 2.1 Subjects

Nine young healthy individuals participated in this study (mean age 31.22; SD 4.55; 5 males). All subjects were consistently right handed, as determined by the Edinburgh Handedness Inventory [40] with a mean laterality quotient of 97. Subjects were recruited from an open access advertisement posted on a website for subject recruitment (www.forsøgsperson.dk). All subjects gave their written informed consent. The experimental protocol (H-18031987) has been approved by the Regional Committee on Health Research Ethics of the Capital Region of Denmark.

### 2.2. Study design

Participants underwent a single pcASL-MRI session that lasted approximately 50 minutes. The experimental procedures are illustrated in Figure 1. Bipolar aTDCS was applied with the anode targeting left primary motor cortex (M1) and the cathode on the right side of the forehead (supraorbital (SO) region). During stimulation, participants were resting in the MRI scanner with their eyes focusing on a fixation cross displayed in the middle of a screen.

**Figure 1:**
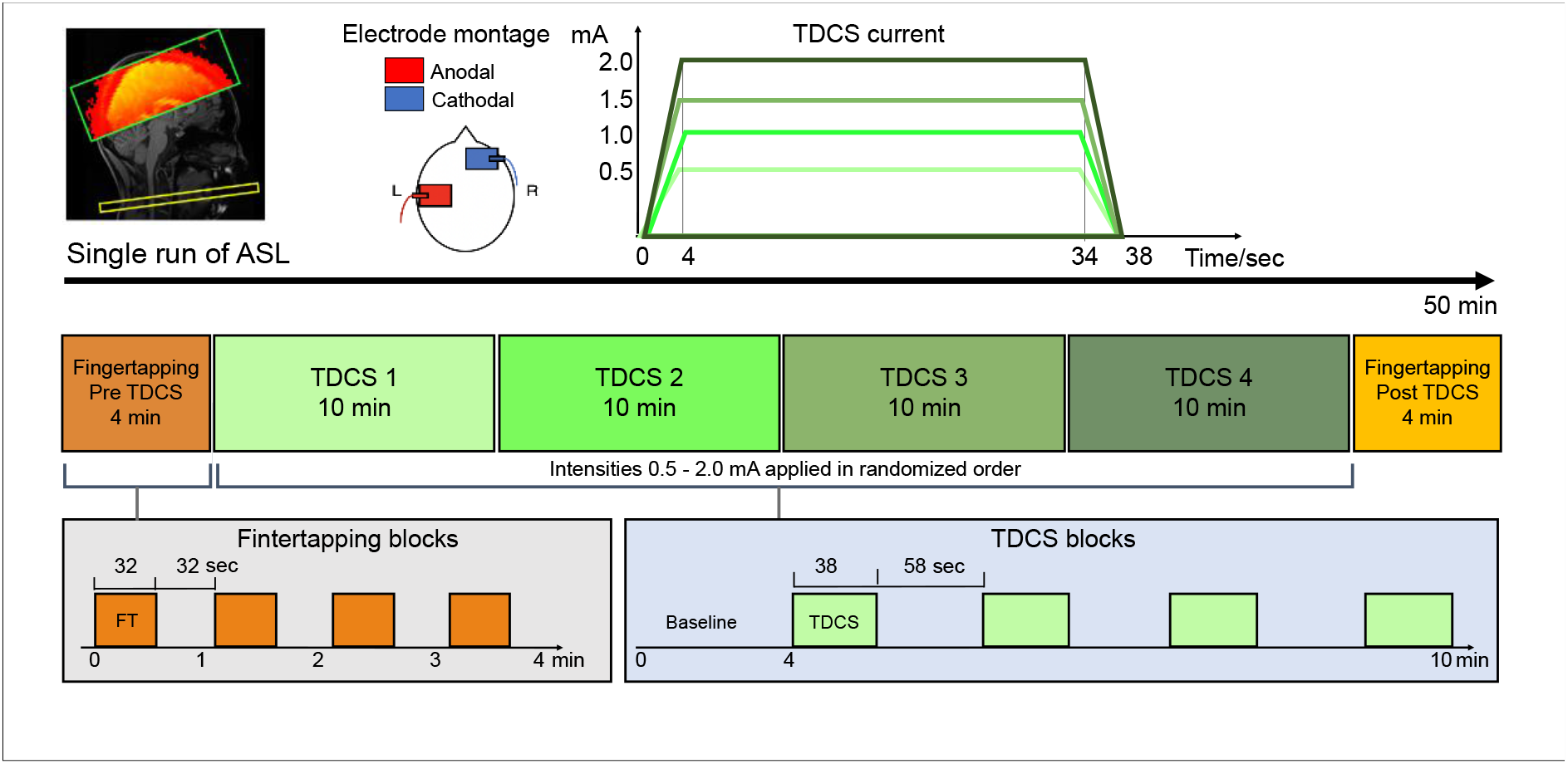
Experimental design Legend text: **Figure 1**. Experimental design. *Top:* tDCS was applied with 5×7cm^2^ square electrodes with anodal on C4 and cathodal on the right supraorbital area. We applied four different current intensities of 0.5, 1.0, 1.5 and 2.0 mA. Current waveform consisted of 4 sec ramp-up, 30 sec stimulation and 4 sec ramp-down. *Middle and bottom:* We performed a single run of pcASL, with a block of 4 min finger-tapping (FT) followed by four consecutive 10 min blocks of stimulation (tDCS) with randomized order of intensities, ending with a block of 4 min FT. Total duration of the scanning session is approximately 50 minutes.

**Figure 2:**
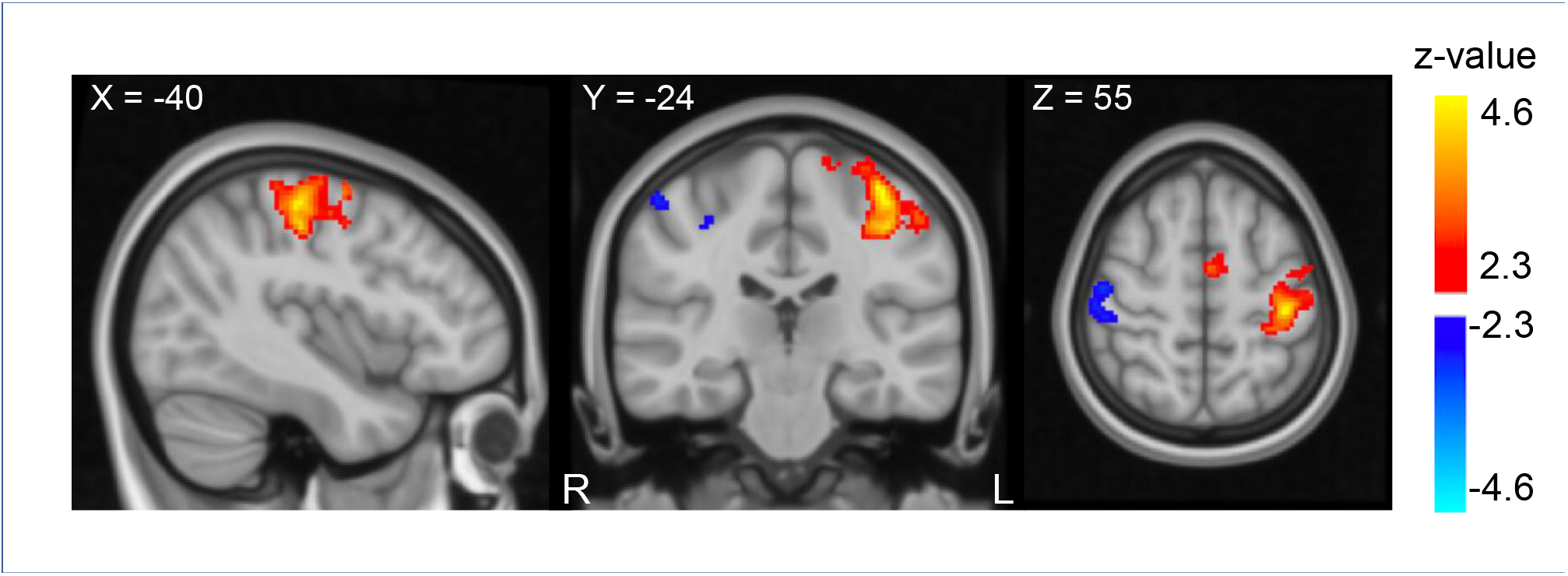
Perfusion activity during finger-tapping Legend text: **Figure 2:** Perfusion changes during right hand finger-tapping before stimulation (FT-pre) on all subjects. Regions show changes in perfusion during FT compared to resting state. Subjects show increase in perfusion corresponding to the left sensorimotor cortex (red), (Cluster size = 1294, Z-max = 4.68, peak coordinate: -38, -24, 60, cluster p-value = 1.72e-08) and decreased perfusion corresponding to the right sensorimotor cortex (blue) (Cluster size = 244, Z-max = 3.0, peak coordinate: 50, -14, 54, cluster p-value = 0.047). Results are overlaid on MNI template. The figure shows the results for a voxel-wise threshold at *p* = 0.01 (Z=2.3) and cluster corrected threshold of p = 0.05 for illustrative purposes. Perfusion changes during tDCS did not survive the cluster-corrected level. The primary analysis was performed with a cluster-defining threshold of *p*= 0.001 (Z=3.1) at the voxel level and cluster corrected threshold of p = 0.05 (shown in supplementary figure 2).

The experiment included four aTDCS blocks. During a TDCS block, one of four current intensities were applied. Target current intensity was set at 0.5 mA, 1.0 mA, 1.5 mA or 2.0 mA in a given block. The order of blocks was pseudorandomized and participants were blinded. Each aTDCS block lasted 10 minutes and consisted of 4 min no-stimulation baseline followed by alternating epochs of aTDCS (38 s) and periods without TDCS (58 s). During an aTDCS epoch, stimulation current was linearly ramped up to the target intensity within 4s, continuously applied at target intensity for 30 s, and then ramped down again within 4s.

After each aTDCS block, the participants answered a series of questions through the MR-speaker system about how they had experienced the preceding aTDCS block, using a six-level Visual Analogue Scale (VAS)(see supplementary table 1).

We included finger-tapping (FT) blocks without stimulation at the beginning (FT-pre) and the end (FT-post) of the pcASL-MRI experiment to compare rCBF changes evoked by aTDCS with rCBF change during voluntary motor activity. During FT blocks participants tapped their right (dominant) index and thumb together paced by a blinking cross (2 Hz). Each FT-block was four min, consisting of interleaved 32 s epochs of FT and 32 s of rest.

### 2.3 Transcranial DC stimulation in the MRI scanner

We applied aTDCS at 0.5, 1.0, 1.5 and 2.0 mA. The center of the anodal electrode corresponding to the C3 location of the 10/20 EEG system [41], with 2-mm thick 7×5 cm rubber electrodes (NeuroConn, Illmenau, Germany). Connector plugs were arranged pointing down towards the ear for the anodal electrode, and horizontally outwards for the cathodal electrode. Electrodes were applied with a thin layer of ten-20 conductive gel (Weaver and Company, Aurora, Colorado, US) and fixated with a net cap. Current was applied by a battery-operated DC-stimulator with MR compatible stimulation cables and filter-boxes (NeuroConn Illmenau, Germany). For safety reasons, the total impedance was kept below 15 kΩ, which included the two 5 kΩ resistors in the cables.

### 2.4 MRI Image acquisition

Images were acquired on a Phillips 3 Tesla MR Achieva scanner (Philips, Best, Netherlands) using a 32-channel head coil. A localizer scan assessed the head-position prior to a structural T1-weighted whole-brain scan using a 3d-TFE multi-shot sequence (TR/TE = 6.0/2.7 ms; flip angle = 8°; FOV= 245 FH 245 AP 208 RL mm^3^; isotropic resolution = 0.85 mm^3^) and a T2-weighted whole-brain scan using a 3D-TFE multi-shot sequence (TR/TE = 2500/265 ms; flip angle = 90°; FOV= 245 FH 245 AP 190 RL mm^3^; isotropic resolution = 0.85 mm^3^).

Pseudo-continuous (pc)ASL was acquired as a single run per block, resulting in six ASL-MRI runs per participants, corresponding to the four aTDCS and two FT blocks per participant (Figure 1). The labeling plane was positioned where the Vertebral and Internal Carotic Artery are parallel (approximately at C2 level) and angled perpendicular to the vessel orientations. pcASL images were acquired with background suppression by two pulses, pulsed continuous labeling, label duration of 1650 ms, and post-label-duration of 1200 ms, using an echoplanar imaging (EPI) readout. Image resolution was 3 × 3 × 4 mm. A single ASL volume consisted of 17 slices with a gap of 0.5 mm, covering the pericentral cortices and adjacent frontoparietal regions of both cerebral hemispheres. Duration for each ASL dynamic was 2 × 4.0 sec, with 78 dynamics per aTDCS block, and 32 dynamics per FT block.

### 2.5 Data analysis

All ASL-MRI data was analyzed using FSL software, Wellcome Centre for Integrative Neuroimaging, University of Oxford. (https://www.win.ox.ac.uk).

Pre-processing of ASL-MRI data is described in Appendix A. ASL-MRI data were analyzed using FSL FEAT. For each participant, we did six separate first level analyses, one for each of the four aTDCS runs, and two for FT-runs. Voxel-wise changes in rCBF were analyzed by fitting a General Linear Model (GLM) to the time series of each ASL-MRI run/block modelling either one of the four aTDCS intensities or the movement sequences. The GLM featured three regressors, as previously described in Moisa et al. [42] (further details describing our GLM in Appendix A). The positive z-activation map from the perfusion regressor was used for group level analyses, described below.

#### 2.5.1 Definition of volumes of interests

The left M1-HAND was our primary volume of interest (VOI) . We also wanted to determine specific functional changes in sub-regions within the primary VOI, based on depth and proximity to the anodal electrode and we planned to include the region with the highest induced E-field as an additional VOI:

##### 2.5.1.1 Functional defined VOI of the left M1-HAND (M1_FT_)

We used the average FT-pre activation as a functional localizer for this VOI (Figure 3A, Figure 4). The VOI was derived from a whole-brain voxel-wise group analysis using a one-sided t-test with a corrected statistical threshold of p<0.05 set for family wise error (FWE) cluster level correction and a cluster-defining threshold that was set to p <0.001 at the voxel level (corresponding to Z=3.1).

**Figure 3:**
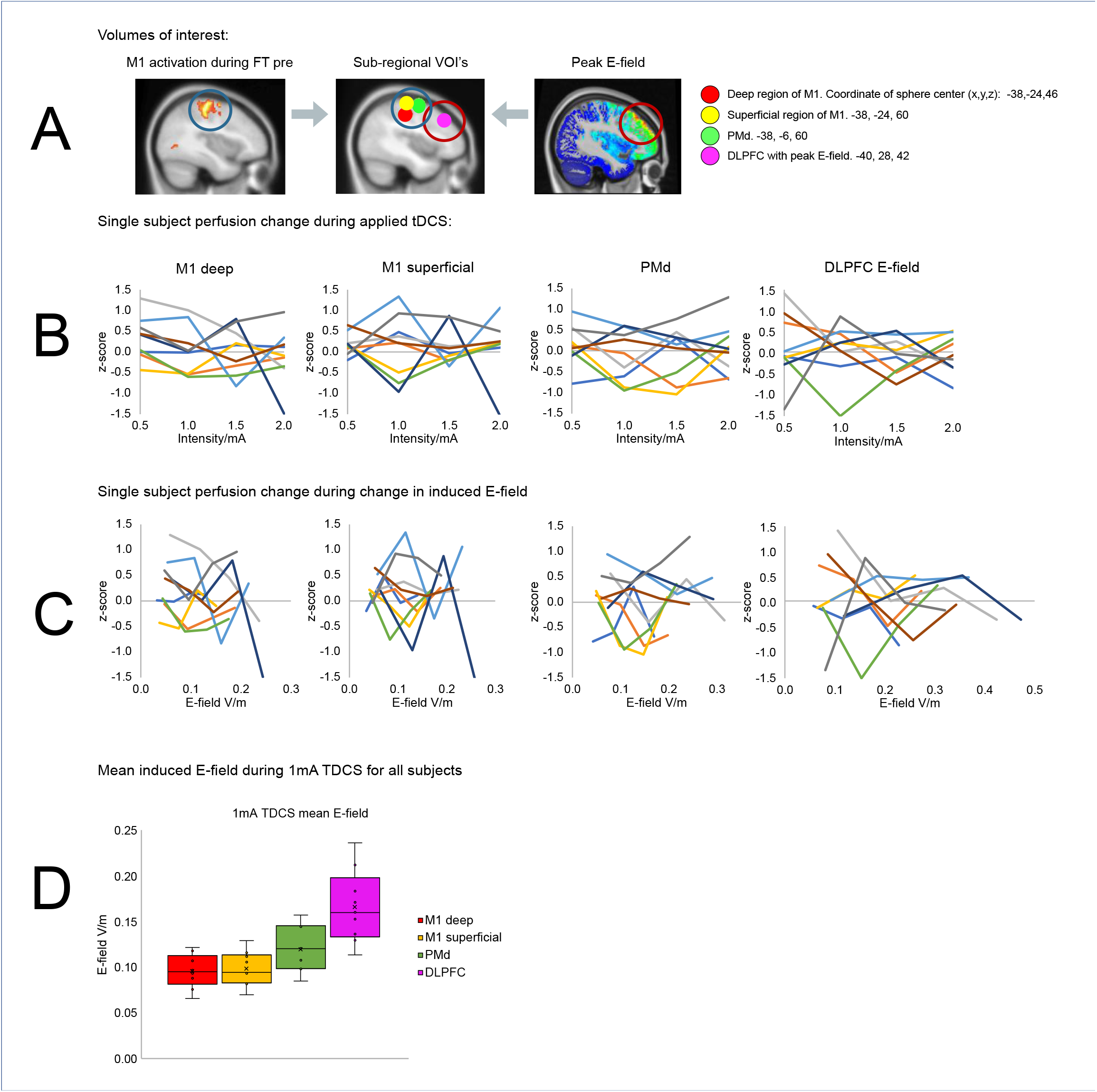
Change in cerebral blood flow within regional volumes of interest Legend text: **Figure 3**. Individual changes in regional cerebral blood flow within four spherical volumes of interest (VoIs) with a radius of 10mm. *Panel A:* The regional peak activations during finger-tapping prior to stimulation (FT-pre) was used as a functional localizer for the three spheres. Spheres were placed in the superficial primary motor cortex (M1) (yellow, center coordinate: -38; -24; 60,) the deep part of M1 (red, center coordinate: -38; -24; 46) and the left dorsal premotor cortex (green, center coordinates -38;-6;60). The fourth VoI (magenta) was placed at the location of maximum induced E-field in all subjects (center coordinate -40;28;42). *Panel B:* Plots show individual activation within each VoI for each stimulation intensity. *Panel C:* Plots show individual activation within each VoI as a function of the induced E-field instead of tDCS intensity. *Panel D:* boxplot of mean induced E-field in all four VOI’s at 1 mA TDCS. Y-axis shows induced E-field V/m. Outliers defined being beyond 1.5 inter-quartile range of each quartile.

**Figure 4:**
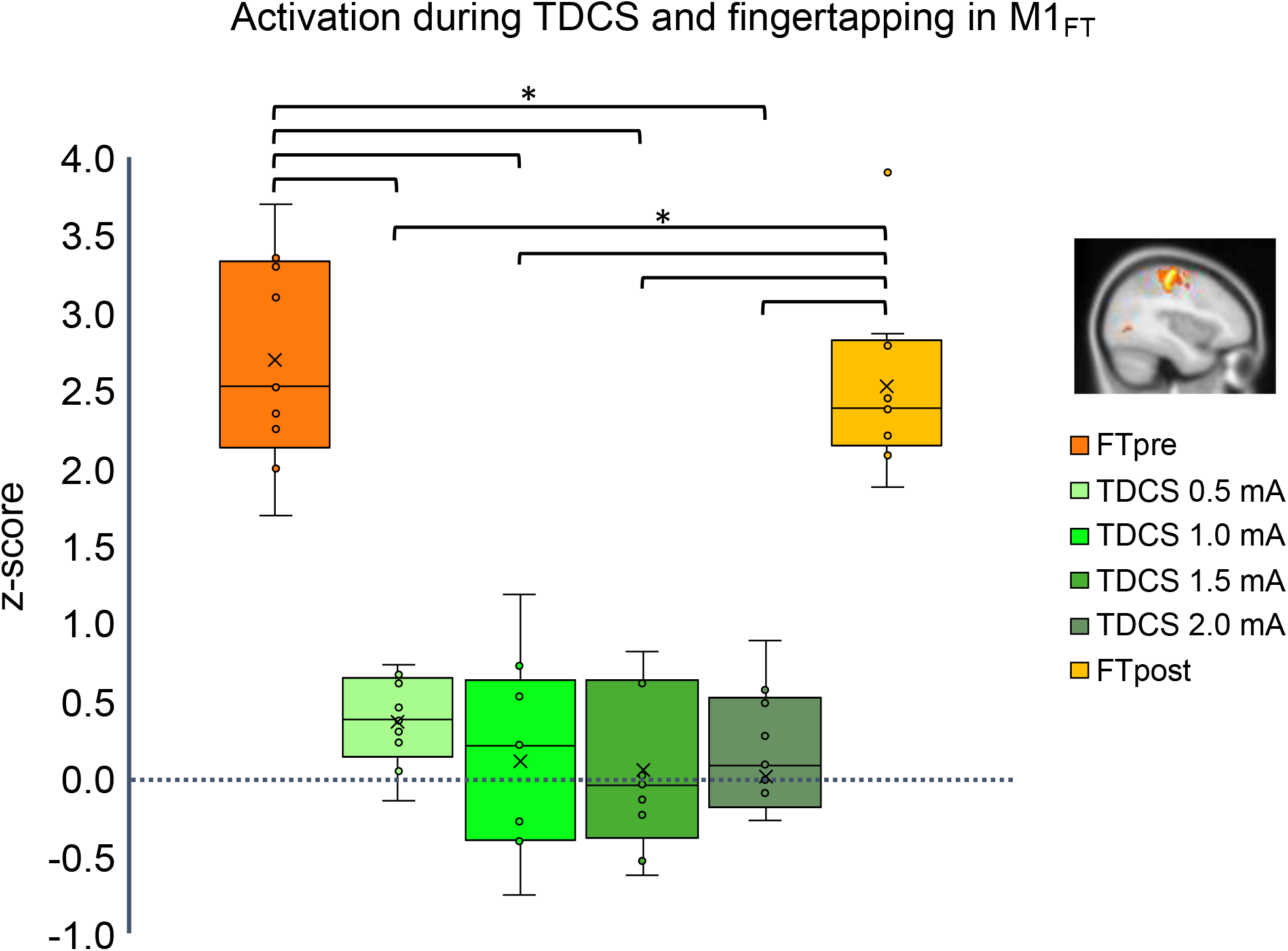
change in regional cerebral blood flow in the left M1_FT_ VOI Legend text: **Figure 4**. Change in regional cerebral blood flow in left primary motor cortex. The region is defined corresponding to the regional activation during the finger-tapping task prior to stimulation (FTpre) as shown red colors in fig 1. Box and whisker plot of activation during tDCS at 0.5 – 2.0 mA (green shades) and finger-tapping before stimulation (FTpre) and after (FTpost) (orange shades). Y-axis shows activation in t-score. Outliers defined being beyond 1.5 inter-quartile range of each quartile.

##### 2.5.1.2 Anatomical defined VOI of the left primary sensorimotor cortex (SM1_anat_)

To avoid bias from the motor-task guided VOI (M1_FT_), we additionally defined an anatomical VOI around the left primary sensory-motor cortex based on the probabilistic atlas from FSL’s Juelich map, including areas BA4a, BA4b BA3a, BA3b and BA6 for the left hemisphere. The probabilistic area masks were thresholded at 50% relative to their peak values and binarized. In addition, the extend of the mask was restricted to fit area around M1-HAND by including only voxels with MNI coordinates between -52 mm and -25 mm in left-right direction and ≥36 mm in inferior-posterior direction (Figure 5).

**Figure 5:**
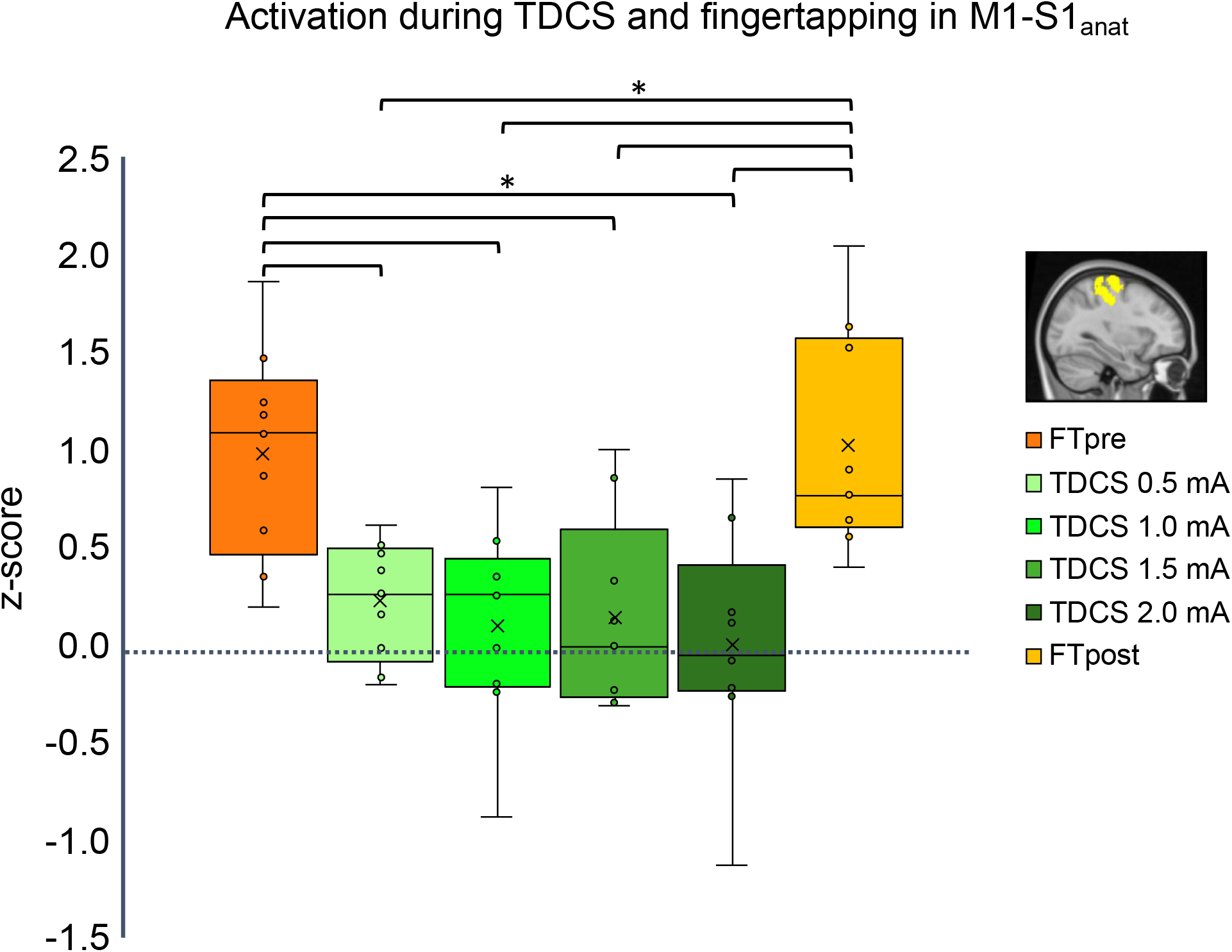
change in regional cerebral blood flow in the left M1-S1_anat_ VOI Legend text: **Figure 5**. Change in regional cerebral blood flow in left primary sensorimotor cortex. The region is defined using the probabilistic anatomical Juelich atlas. Areas BA4a, BA4b BA3a, BA3b and BA6 for the left hemisphere was used, and the mask was limited in each direction to fit the hand knob. Box and whisker plot of activation during tDCS at 0.5 – 2.0 mA (green shades) and finger-tapping before stimulation (FTpre) and after (FTpost) (orange shades). Y-axis shows activation in t-score. Outliers defined being beyond 1.5 inter-quartile range of each quartile.

##### 2.5.1.3. Sub-regional spherical VOI’s in precentral cortex

Secondary VOI analyses included three precentral sub-regions and the frontal cortical site that was exposed to the maximal electrical field during aTDCS. We defined three VOIs within the left pre-central gyrus (10 mm spheres) using the FT-pre as a functional localizer. The VOIs were placed at the regional activity peaks within M1_FT_, (Figure 3A). Two of the VOIs were placed at the superficial and deep point of highest regional activation in the primary motor cortex, “M_deep_” and “M1_superficial_” (center coordinates x,y,z: [-38, -24, 46] and [-38, -24, 60]). The third VOI was placed in a functional activated area corresponding to area 6 in the Juelich atlas in FSL, the dorsal premotor cortex “PMd” (center coordinate [-38, -6, 60]).

We performed additional analysis that considered the current distribution in each subject. We used individual structural scans to simulate the induced electric field for each subject, using SimNIBS v.3.2.2 [21]. For details on E-field simulations, please see Appendix A (and Figure 3). To define a VOI for the maximal electrical field each individual E-field simulation at 1 mA were overlaid in MNI-space, considering only positions at which the gray matter of a least 5 subjects overlapped. A 10mm VOI was placed at the gray matter location at maximum of the averaged E-field (sphere center coordinate: -40, 28, 42), corresponding to the left dorsolateral prefrontal cortex (DLPFC) (bottom right panel in supplementary S3).

#### 2.5.2 Statistical inference on group level

##### 2.5.2.1 Regional perfusion changes in the cortical VOIs

We ran six GLM analyses at group level, one for each aTDCS condition, corresponding to the four aTDCS intensities, and the two finger tapping sessions. The parameter estimates from the perfusion regressor from the first level analyses was fitted into a second level GLM that modelled the group average, corresponding to a one-sided t-test for each condition. In each model, the z-scores were averaged within our pre-defined volumes of interest (VOI definitions described in section 2.5.3). For each VOI, we submitted the spatially averaged z-scores into a two-way repeated measure ANOVA for within subject factor “conditions” with 6 levels (4 aTDCS intensities, 2 movement sequences). We performed post-hoc tests with pairwise comparisons, p-values were corrected for multiple comparisons using the Bonferroni method. For all ANOVA’s, we used Mauchly’s test to test for sphericity, and corrected with the Greenhouse Geisser method when sphericity was violated. ANOVA and post hoc tests were done using version 25 of the SPSS statistics software package (IBM, Armonk, New York, USA).

##### 2.5.2.2 Correlation between regional E-field and regional perfusion change

We explored the relation between the highest induced E-field and perfusion response on an individual level. Here we simulated the E-field (at 0.5, 1.0, 1.5, 2.0 mA) in all four VOIs (M1_deep_, M1_superficial_, PMd, DLPFC_E-field_) in all subjects. For each intensity and each VOI, we ran linear regression analyses on the mean perfusion activation and E-field, to see if there were any correlation between the individual perfusion changes and estimated individual E-field strength (see supplementary S4). We used Bonferroni correction between the 16 comparisons, with an alpha level of 0.05.

##### 2.5.2.3 Voxel-based exploratory analyses

We performed a complementary voxel-based analysis to test for linear increase or decrease in perfusion throughout all aTDCS intensities, including all voxels. We applied a cluster-size threshold of p = 0.05 and voxel-wise threshold of z = 3.1 or p = 0.001. We additionally explored finding potential activity with a more liberal voxel-wise threshold of z = 2.3 or p = 0.01.

##### 2.5.4.4 Analysis of psychometric data

For details on analysis of VAS-scale rated sensory experience, please see Appendix A.

## Results

None of the participants experienced major side effects during the study. The results of the psychometric assessment are presented in Appendix A as supplementary material.

### 3.1 Regional perfusion changes in the precentral M1-HAND target region

We first examined mean perfusion changes during FT and during aTDCS in the M1_FT_ VOI. We found a significant main effect of “conditions”, indicating a difference in mean rCBF within M1_FT_ among the experimental conditions (F = 54.205, p < 0.000). Pairwise comparisons showed a significantly higher rCBF during FT compared to all aTDCS intensities after Bonferroni correction (Table 1). FT induced rCBF increases after the aTDCS blocks did not differ from FT induced perfusion prior to aTDCS (FTpost vs. FTpre). There was no significant difference in rCBF between any of the applied aTDCS intensities, but mean rCBF during 0.5mA aTDCS was slightly higher than baseline rCBF (Figure 4). Post-hoc exploratory one-sample t-tests on each intensity condition revealed an increased perfusion (z-score) in M1_FT_ at 0.5 mA aTDCS compared to baseline, p_uncorrected_ = 0.005 which survived Bonferroni correction (p_Bonferroni_ = 0.02).

**Table 1.** Pairwise comparisons between fingertapping and TDCS activation (M1 FT)

|  | FTpre | FTpost | 0.5 mA | 1.0 mA | 1.5 mA | 2.0 mA |
| --- | --- | --- | --- | --- | --- | --- |
| FTpre |  | 0.487 | 0.000* | 0.000* | 0.000* | 0.000* |
| FTpost |  |  | 0.000* | 0.000* | 0.000* | 0.000* |
| 0.5 mA |  |  |  | 0.215 | 0.128 | 0.256 |
| 1.0 mA |  |  |  |  | 0.837 | 0.650 |
| 1.5 mA |  |  |  |  |  | 0.917 |
| 2.0 mA |  |  |  |  |  |  |

\* significance level after Bonferroni correction at $p \leq 0.003$ (alpha = 0.05)

To control for any potential bias introduced by using a group-specific functional VOI (M1_FT_), we also analyzed rCBF changes using an anatomically defined VOI (Figure 5).

The rCBF changes in the M1-S1_anat_VOI were less strong compared to M1_FT_, but revealed a similar response pattern. Two-way repeated measures ANOVA revealed a significant main effect of “conditions” (F = 9.436, p < 0.000). Pairwise comparisons showed significant difference between rCBF during FT compared to all stimulation intensities after Bonferroni correction (Table 2), with no difference between FTpost vs. FTpre and no significant difference in rCBF between the aTDCS intensities. One-sample t-test on each aTDCS intensity also revealed increased perfusion at 0.5mA compared to baseline, p_uncorrected_ = 0.028, but the increase in rCBF during aTDCS at 0.5mA did not survive correction for multiple comparisons (p_Bonferroni_ = 0.112).

**Table 2.**
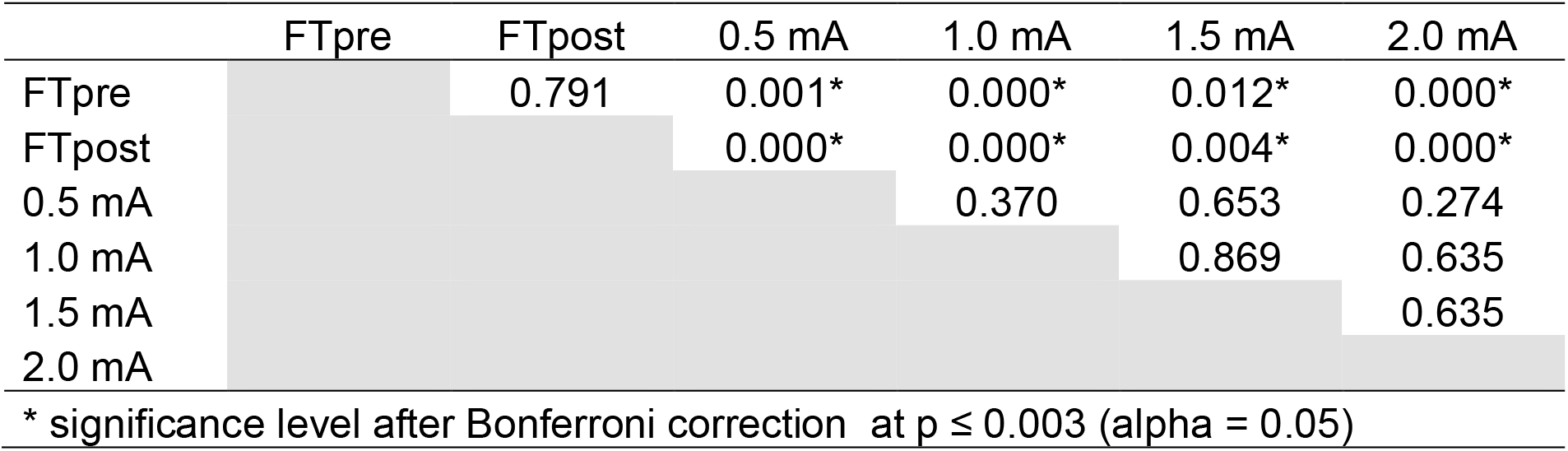
Pairwise comparisons between fingertapping and TDCS activation (M1-S1 anat)

|  | FTpre | FTpost | 0.5 mA | 1.0 mA | 1.5 mA | 2.0 mA |
| --- | --- | --- | --- | --- | --- | --- |
| FTpre |  | 0.791 | 0.001* | 0.000* | 0.012* | 0.000* |
| FTpost |  |  | 0.000* | 0.000* | 0.004* | 0.000* |
| 0.5 mA |  |  |  | 0.370 | 0.653 | 0.274 |
| 1.0 mA |  |  |  |  | 0.869 | 0.635 |
| 1.5 mA |  |  |  |  |  | 0.635 |
| 2.0 mA |  |  |  |  |  |  |

\* significance level after Bonferroni correction at $p \leq 0.003$ (alpha = 0.05)

**Table 3.** Experience of tingling compared between TDCS intensies

|  | 0.5 mA |  | 1.0 mA |  | 1.5 mA |  | 2.0 mA |  |
| --- | --- | --- | --- | --- | --- | --- | --- | --- |
|  | Z-score | p-value | Z-score | p-value | Z-score | p-value | Z-score | p-value |
| 0.5 mA |  |  | -2.000 | 0.046* | -1.604 | 0.109 | -2.041 | 0.041* |
| 1.0 mA |  |  |  |  | -0.333 | 0.739 | -1.604 | 0.109 |
| 1.5 mA |  |  |  |  |  |  | -1.414 | 0.157 |
| 2.0 mA |  |  |  |  |  |  |  |  |

\* significance level after Bonferroni correction at $p \leq 0.008$ (alpha = 0.05)

Analysis of aTDCS-related perfusion changes in the sub-regions *M1*_*deep*_, *M1*_*superficial*_ and *PMd* revealed no significant main effect of aTDCS intensity on rCBF in any of the functionally defined sub-regional VOIs (VOI_M1deep_: F=1.04 p = 0.38, VOI_M1superficial_: F= 0.33 p = 0.81, VOI_PMd_: F=0.63 p = 0.61) (Figure 3)

### 3.2. Perfusion changes in the cortical region exposed to the highest E-field

The aTDCS-induced electrical field reached its maximum in the dorsolateral prefrontal cortex (DLPFC_E-field_) (Figure 3A). The mean E-field in the dorsolateral prefrontal VOI (*DLPFC*_*E-field*_) was consistently higher than the mean E-field in the precentral VOIs (Figure 3D). We did not find any significant effect of aTDCS or FT on mean rCBF in the DLPFC_E-field_ VOI (F = 0.290 p = 0.916) (Figure 3B), individual rCBF activity in the E-field guided prefrontal VOI was highly variable, showing no consistent intensity dependent response pattern.

### 3.3 Whole-brain analysis

Whole-brain analysis revealed no significant increase in rCBF for any of the four aTDCS conditions, or any linear voxel-wise increase or decrease. This was different for the finger tapping task. Right-hand FT before aTDCS (FTpre) lead to increase in rCBF in left M1-HAND (Figure 2 and Supplementary Figure 2). Exploratory analysis at a more liberal cluster-forming threshold (z = 2.3 or p = 0.01) also showed a tapping-associated decrease of rCBF in the right primary sensorimotor cortex ipsilateral to the tapping right hand (Figure 2). FT after stimulation (FTpost) with the right hand also led to increase in rCBF in left M1-HAND (Supplementary Figure 2).

### 3.4 Relationship between induced E-field and regional perfusion changes

Neither for M1_superficial_, M1_deep_, PMd and or DLPFC_E-field_, we found a consistent relationship between regional perfusion changes and the induced E-field in our participants (Figure 3C). We tested whether the perfusion levels in these VOIs could be predicted by individual induced E-field by specifying a linear regression model to predict perfusion activity (z-score) from E-field at each of the four aTDCS intensities. Only the E-fields induced with aTDCS at an intensity of 0.5 mA significantly predicted z-score in M1_deep_, F = 9.333, p_Uncorrected_ = 0.018, R^2^ = 0.571. This however did not survive Bonferroni correction for multiple comparisons (p_Bonferroni_= 0.288). The corresponding plots are shown in Supplementary Figure 4.

## Discussion

Using pcASL-MRI, we measured relative changes in regional cerebral blood flow as proxy for regional neural activity during short epochs of bipolar aTDCS targeting the left M1-HAND. In different aTDCS blocks, we varied the intensity of aTDCS, applying currents at 0.5, 1.0. 1.5, and 2.0 mA. We found no evidence in support of the hypothesis that aTDCS produces intensity-dependent increases in rCBF in the cortical target region. This was the case for all subregions of the targeted pericentral cortex but also in the portion of the laterodorsal prefrontal cortex where the induced E-field was highest.

### 4.1 No modulation of regional blood flow in M1-HAND during short-duration aTDCS

We found no change in the regional perfusion level during aTDCS compared to baseline. This negative finding is in apparent contrast to previous ASL-MRI studies that reported a regional increase in cortical perfusion underneath the electrode [29,35]. Since previous studies used markedly longer stimulation blocks, lasting 8-20 min, it is likely that prolonged TDCS protocols may result in more pronounced effects on rCBF. The effects on rCBF produced by longer TDCS protocols may not reflect instantaneous effects that are directly related to aTDCS but rather secondary neuromodulatory effects of TDCS [43]. Since the neuromodulatory effects of TDCS are longer lasting, they may not be rapidly reversible after the termination of TDCS [43,44]. Hence, it would not be feasible to use longer protocols, if the goal is to determine the individual dose-response profile to aTDCS for personalized dosage control.

The absence of reliable rCBF changes in response to 30-second periods of aTDCS raises the question whether short protocols are sufficient to shift corticomotor excitability in the targeted M1-HAND at all. Here it is important to note that TMS of M1-HAND did reveal a significant increase in cortical excitability already after a short period (i.e., 4 seconds) of low intensity (1mA) aTDCS [13]. As we did not find a comparable immediate increase in regional perfusion, we hypothesize that the rCBF response probed with ASL is less responsive to the polarization effects of TDCS than the pyramidal cortico-spinal neurons, that are trans-synaptically probed with TMS.

### 4.2 Robust rCBF increase in left M1-HAND during finger tapping

While we were not able to detect stimulation-related rCBF changes, M1-HAND showed a robust motor-task induced increase in rCBF during contralateral finger tapping. This indicates that our pcASL measurements were sensitive to increases in regional neuronal activity. When using a more liberal threshold, there was also a trend towards a decrease in regional perfusion in the ipsilateral (right) primary sensory cortex. The task-induced ipsilateral perfusion decrease has not been studied with ASL previously but supports previous neurophysiological findings on interhemispheric interaction [45], and could be utilized by future studies to investigate disruptions of the interhemispheric network during conditions like stroke. Together, the demonstration of task-related bi-directional changes in M1-HAND activity suggests that the lack of any stimulation-induced effects on rCBF in M1-HAND cannot be attributed to an inability of our ASL sequence to detect perfusion changes. Comparing the perfusion during finger tapping before and after the stimulation blocks suggested that the four aTDCS blocks did not induce a change in rCBF that outlasted stimulation. This is consistent with previous BOLD fMRI studies, showing no significant change of task-related activation after consecutive blocks of 20 sec TDCS [46] and 5 min TDCS [26].

### 4.3 No dose-dependent changes in regional blood flow during short-duration TDCS

We did not find any intensity-related difference in rCBF levels during short-duration aTDCS. Previous studies have suggested a non-linear relationship between changes in neuronal activity and the current intensity of TDCS, [47,48] with lower currents potentially inducing stronger effects than high currents [49]. Of note, we found a small but consistent effect of low-intensity aTDCS at 0.5 mA on rCBF in the deep region of M1-HAND (Figure 4). This activity-enhancing effect at the lowest current intensity may reflect stochastic resonance effects which may only emerge at very weak electric fields [50].

We did not apply aTDCS at intensities above 2mA. Therefore, it is possible that higher current intensities might be needed for causing a consistent change in regional perfusion with short-duration aTDCS. This hypothesis is supported by recent findings from Jonker et al. who did not find any effect on cortical excitability after 20 minutes of aTDCS at 2mA [17] together with in vivo measurements of neural spiking activity in animals [20] and humans [19][51]. These measurements suggest that convential TDCS at 2mA current induces electrical field gradients at the lower end of needed to induce neuronal spiking activity. Our negative results thus emphasizes the relevance of exploring the effect of TDCS at higher current intensities in future experiments.

### 4.4 Difference in target engagement in sub-regional VOIs

Recent biophysical models suggest that the effect of the induced current is dependent on the orientation of the targeted neurons. Pyramidal tract neurons have lower thresholds for activation at the axon terminal and node of Ranvier [52], and the depolarization of these segments is highly dependent on their orientation to the electric field: Neurons in the gyrus crown are oriented perpendicular to the E-field and depolarization will predominantly occur at the proximal part of the axon [52,53], whereas neurons located in the sulcal depth will not achieve the same level of polarization. Different susceptibility to electric stimulation in superficial versus deeper structures is supported by E-field simulations showing that the induced electrical field is highest at the gyral crown [54]. We therefore expected higher engagement of aTDCS in the superficial M1. Contrary to our hypothesis, aTDCS did not induce regional-specific perfusion changes, neither in the superficial M1-HAND nor in deeper regions or the adjacent PMd. The inability to detect any region-specific effects on rCBF might be attributed to the low current intensities and the use of a classical non-focal electrode montage.

### 4.5 E-field simulations of target engagement

Our E-field simulations revealed that bipolar aTDCS induced its maximum E-field more rostrally in the frontal cortex than the intended stimulation target (M1). There was however no significant aTDCS induced change in rCBF in the area of maximal current density either. At the group level, the peak E-field location was located in the left dorsolateral prefrontal cortex. This finding is consistent with previous work showing that the maximum TDCS-induced E-field is not necessarily located directly underneath the stimulation electrodes [55]. In a recent ASL-TDCS study, Jamil et al. reported that 15 minutes of bipolar TDCS targeting M1, induced polarity-specific changes in rCBF under the M1-electrode, together with regional E-field dose-dependency with the strongest correlation at TDCS intensities between 1 – 2 mA [37]. In our study, we used the same montage and intensity range, but applied short period TDCS. The short periods of TDCS showed no correlation between the regional induced E-fields and rCBF. Interestingly, the individually induced E-field in the deep region of M1-HAND predicted the rCBF increase in the deep region of M1-HAND during low-intensity aTDCS at 0.5 mA. This finding suggests that for short lasting TDCS, the induced E-field scales with the neuronal activation in the deeper part of M1-HAND in a dose-dependent manner only at low current strength. Stochastic resonance may explain why variations within an apparently narrow range of weak currents positively correlate with the regional neural response [50,56].

In summary, We consistently found that the immediate perfusion responses to short-duration aTDCS are highly variable among subjects (as presented in Figure 4). On the one hand, this inter-individual variability challenges the common practice to apply a fixed stimulation intensity in TDCS studies, as this will most likely evoke substantially different physiological responses across individuals. On the other hand, the fact that we could not find a clear E-field-dependent change in regional CBF, indicates that our setup may not be a feasible method to define individual dose-response relationships and to guide the personalized dosing of TDCS.

### 4.6 Limitations

We would like to mention some limitations of our study.

We used pcASL-MRI to measure changes in regional perfusion as proxy read-out of regional neural activity. Since the ASL signal is noisy, we may have missed subtle changes in regional perfusion. Since our pcASL MRI approach reliably detected movement related activation of M1-HAND in each individual, we argue that our approach was sensitive enough to detect task-related fluctuations in regional neuronal activity. We were also able to detect a subtle increase in rCBF at the lowest stimulus intensity (aTDCS at 0.5 mA), suggesting that the pcASL MRI approach was sufficiently sensitive with respect to the experimental question.

A recent review emphasized that TDCS may produce direct vascular responses [57], including an immediate vasodilatory effect. Therefore, one has to consider direct vascular effects in case of TDCS related changes in cortical perfusion. Since we found no consistent dose-dependent increases in rCBF, we argue that short-duration TDCS neither produced dose-dependent neurovascular nor primary vascular effects in the M1-HAND. Since movement related activation was unchanged after the atDCS blocks, we also infer that the short-duration blocks did not produce substantial after effects on task-related neurovascular coupling in M1-HAND.

The experimental design did not include a sham condition to control for off-target effects of aTDCS for instance due to somatosensory co-stimulation. We also did not test for polarity dependent effects, as we only targeted the M1-HAND with anodal stimulation. However, as we have not found any significant changes in regional cortical perfusion related to stimulation, we argue that this does not reduce the reliability of the results. Furthermore, the group size was small, but sufficiently large to capture the individual dose-response relation and to demonstrate substantial inter-individual variability. Future studies that are designed to provide more insight into target engagement and personalization of TDCS may focus more on focal electrode montages [58], taking individual anatomy into consideration for electrode placement [54], and investigating the tolerability and effect of higher current intensities [19,20].

## Conclusion

We show that pcASL does not reveal consistent immediate effects of short-duration aTDCS on neural activity (as reflected by regional perfusion), but that reliably picks up activity changes during a simple motor task, by catching perfusion increase in the contralateral motor cortex and simultaneous ipsilateral perfusion decrease. This discrepancy between the lack of reliable aTDCS activation and the presence of a clear movement-related activation, shows that the immediate aTDCS effects on regional neural activity during short periods of stimulation are subtle. Longer administration periods may be required to produce more substantial shifts in regional activity, that lead to clear changes in rCBF. Yet such changes might have a primary neural or vascular origin, given that TDCS has primary effects on both the neural and vascular structures of the brain [57].

