## Appendix A for "Short periods of bipolar anodal TDCS induce no instantaneous dose-dependent increase in cerebral blood flow in the targeted human motor cortex"

Methods:

**Pre-processing of ASL-MRI data**

Data were first corrected for motion using FSL mcflirt, followed by non-linear warping (i.e. “normalization”) to the MNI T1 template. To achieve accurate normalization, the control EPI without background suppression acquired before the ASL images (M0 image) was affinely registered to the individual T1 structural image (FSL flirt), and the T1 structural image was non-linearly registered to the MNI T1 template (FSL fnirt). The combined registration steps were then applied to the motion corrected ASL data. We used the M0 image, as the ASL images were acquired with background suppression and had too little contrast to allow for a reliable registration to the T1 image. The motion corrected and normalized ASL images were spatially smoothed with a Gaussian kernel of 8 mm full width half maximum and temporally high pass filtered with a cutoff of 100 seconds.

**First level analysis with GLM**

ASL-MRI data were analyzed using FSL FEAT. For each participant, we did six separate first level analyses, one for each of the four TDCS runs and two movement runs. Voxel-wise changes in rCBF were analyzed by fitting a General Linear Model (GLM) to the time series of each ASL-MRI run/block modelling either one of the four TDCS intensities or the movement sequences. The GLM featured three regressors. The first regressor modelled perfusion baseline, by alternating intensity variation between control and tag images. The second regressor modelled the BOLD activation, with different block regressors for the TDCS and movement runs. For the TDCS runs, the second block regressor modelled the alternation between when stimulation was on and off (initial 4 min baseline followed by interleaved 38 seconds ON, 58 seconds OFF). For the movement runs, the second block regressor modelled alternation between finger-tapping and rest (initial 32 seconds fingertapping followed by 32 seconds rest). The second regressor for the BOLD activation was then modelled as the convolution of the block regressor with a standard hemodynamic response function (HRF). The third regressor modelled the perfusion change, by multiplying the regressor for perfusion baseline (tag-control) and TDCS- or movement-induced BOLD activation.

**E-field simulations:**

We used the T1 and T2 scans to simulate the individually induced electric field, at each applied intensities for the M1-SO montage, using SimNIBS v.3.2.2 [21]. For the simulation, we indicated the electrode montage as described in section 2.3, with a conductive gel thickness of 4mm. The anodal electrode was placed at C3 in the integrated 10/20 EEG-system in SimNIBS (corresponding to left M1), and the cathodal electrode was placed at the right supraorbital ridge of each individual structural reconstruction, guided by Fp2 in the EEG-system (Supplementary figure 3 illustrates each individual simulation at 1mA).

**Analysis of psychometric data:**

All participants were asked to rate their sensory experience of each stimulation intensity on a VAS-scale of 6 levels, ranging from “not at all” to “very much” (see supplementary table 1). We assigned these in an ordinal scale from score 0 – 5 for analysis with Friedman test, with intensity as factors (0.5, 1.0, 1.5 and 2.0 mA), and post-hoc test with Wilcoxon signed rank tests, comparing each intensity against each other. We used Bonferroni correction between the six comparisons, with alpha level of 0.05.

Results:

**Psychometric assessment**

*Tingling/prickling sensation:* Friedman test revealed a significant difference in experience of tingling/prickling sensation on the scalp depending on TDCS stimulation intensity, $X^2(3) = 8.143$, $p = 0.043$. However, post-hoc pairwise comparison with Wilcoxon Signed Ranks test did not show any significant difference between any intensities after Bonferroni correction (table 3). Supplementary figure 5 show a linear increase in prickling/tingling sensation following the increase in current intensity. Concurrent ratings of discomfort were minimal at all intensities.

*Malaise and fatigue:* Participants did not report differences in experience of malaise depending on stimulation intensity ($X^2(3) = 3.585$, $p = 0.310$) nor was there any significant difference in experience of fatigue ($X^2(3) = 0.882$, $p = 0.830$).

*Phosphenes:* Two subjects experienced phosphenes during TDCS. One reported phosphenes during 0.5mA (intensity at VAS-scale 1), the other subject reported during 0.5 (VAS = 1), 1.0 (VAS = 1), and 2.0 mA (VAS = 2). No discomfort was associated with the phosphenes.

*Metal taste:* Two subjects experienced metal taste during stimulation. One subject reported metal taste during all four intensities (VAS = 2 for all). The other subject experienced metal taste during 1.0 and 2.0 mA (VAS = 1 for both). No discomfort was reported in relation to experiences the metal taste.

*Nausea:* None of the subjects experienced nausea.
