## Supplementary figures for "Short periods of bipolar anodal TDCS induce no instantaneous dose-dependent increase in cerebral blood flow in the targeted human motor cortex"

Supplementary material

Supplementary figure 1:

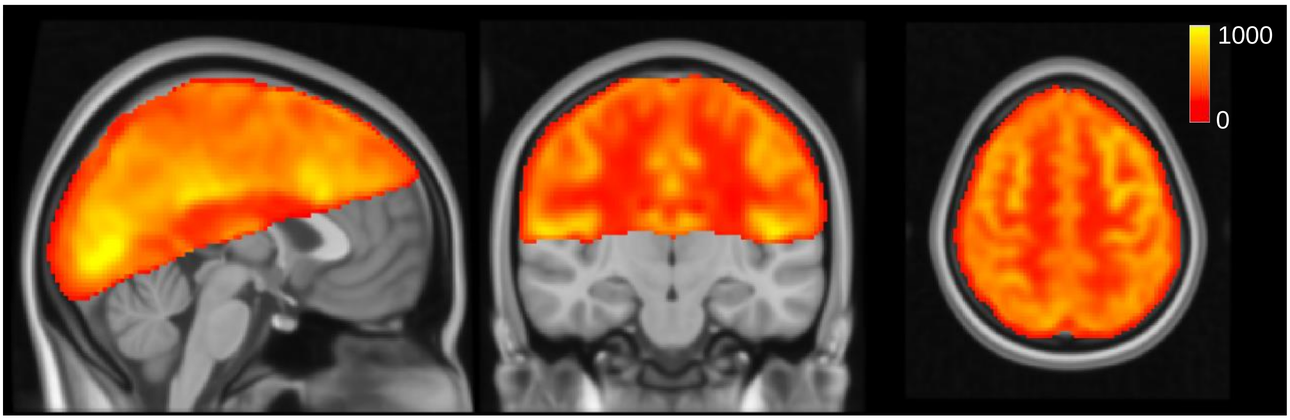


**Supplementary figure 1:** Average perfusion map in arbitrary units for an example subject

Supplementary figure 2:

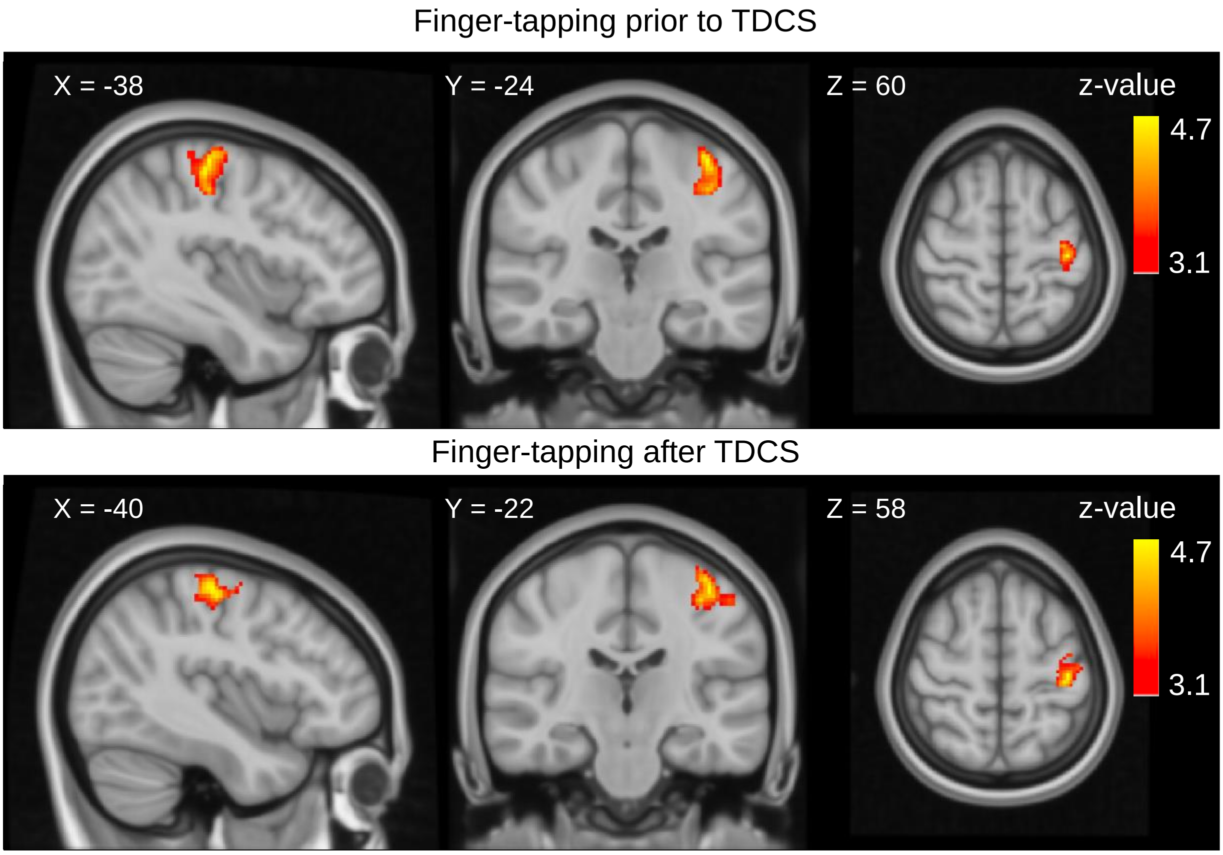


**Supplementary figure 2:** Whole brain analyses shows activation in M1-HAND after finger-tapping. *Top panel* shows grouped activation from all subjects during the right hand finger-tapping task before TDCS (FTpre) (cluster size = 325 voxels, Z-max = 4.68 peak coordinate: -38, -24, 60). *Bottom panel* shows activation after TDCS (FTpost) (cluster size = 445 voxels, Z-max = 4.69 peak coordinate: -40, -22, 58, cluster p-value = 5.96e-08). Coordinates are placed at the global maximum. Voxel-level threshold is defined at $p$= 0.001 (Z=3.1), and cluster corrected threshold of p = 0.05.

Supplementary figure 3:

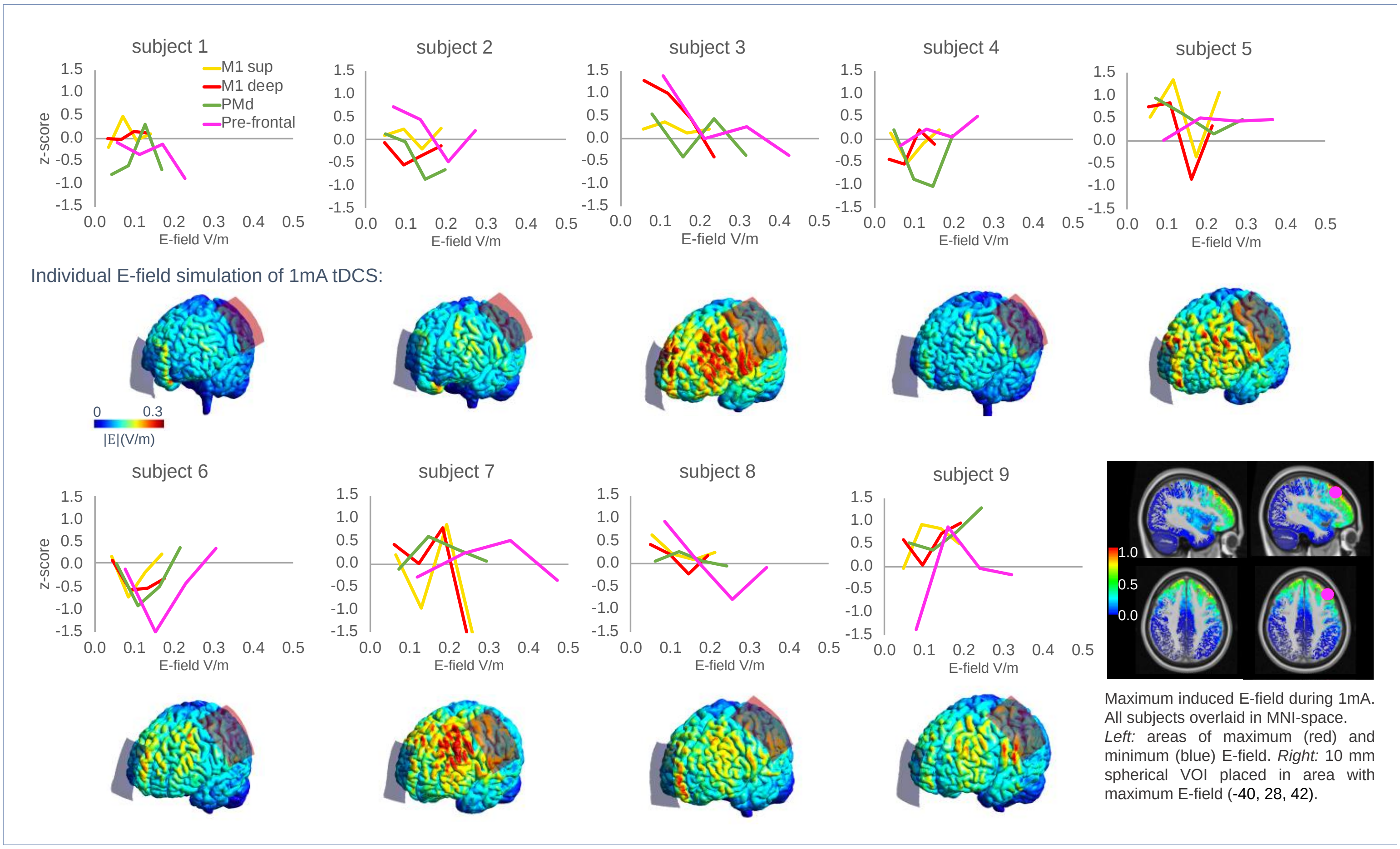


**Supplementary figure 3:**

Individual plots of regional activation and induced E-field. Line plots show individual perfusion change as a function of induced E-field in the four VoIs. For each subject is visualized the individual electric field strength (norm or length of the electric field vectors) at 1mA tDCS for the M1-supraorbital (SO) montage. Strong fields are located at the edge of the electrodes as well as in between the electrodes at pre-frontal areas instead of directly underneath. *Right-bottom corner:* Shows areas of maximum induced E-field at 1mA tDCS, all subjects overlaid on MNI space. Magenta colored VoI is placed at the region of highest E-field.

Supplementary figure 4: Correlation between perfusion response and E-field in subregional regions of interest.

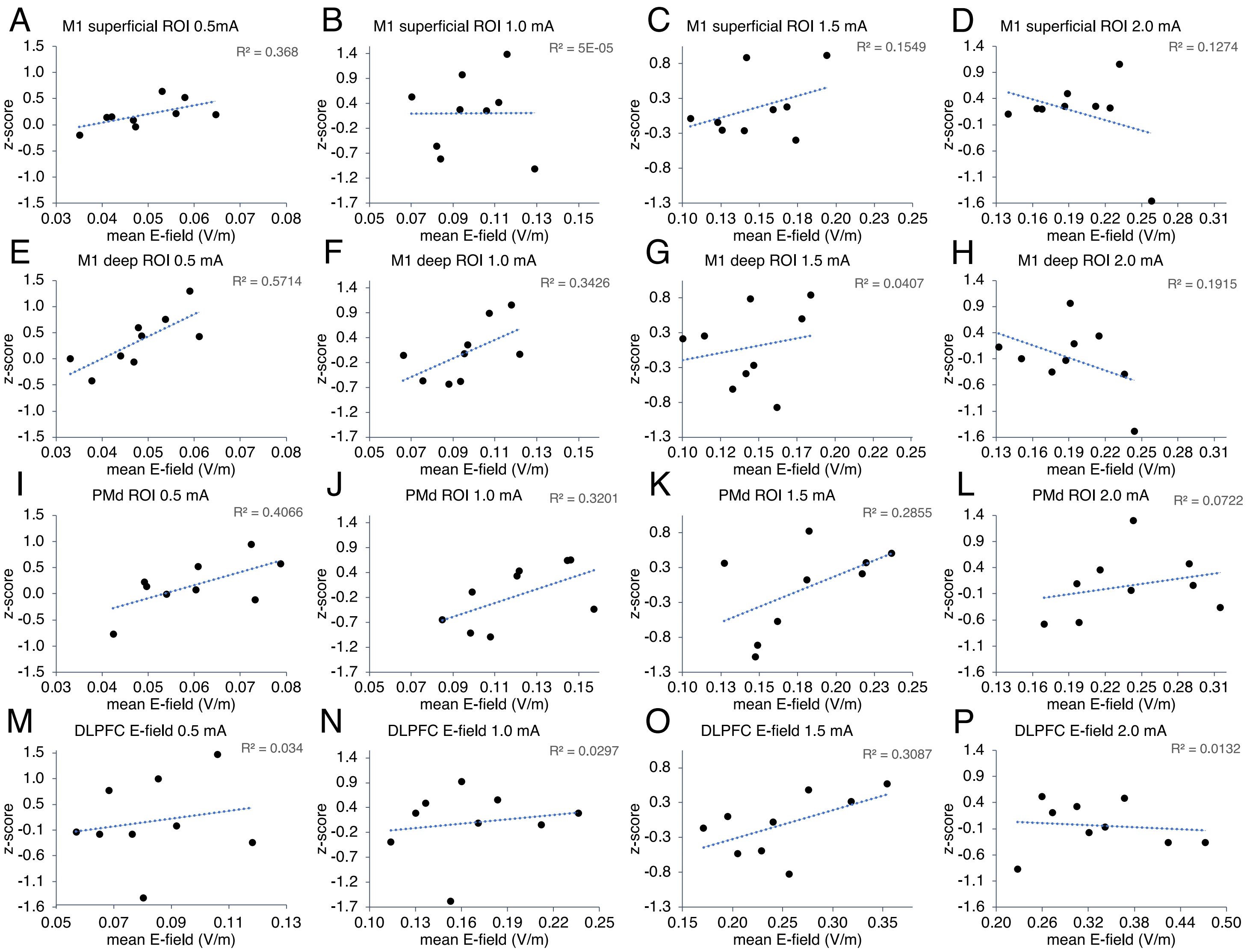


**Supplementary figure 4:** Linear regression plots of activation (t-score) as function of the E-field in the subregional VOI's $M1_{superficial}$ *(A-D)*, $M1_{deep}$*(E-H)*, PMd*(I-L)* and E-field guided VOI $DLPFC_{E\text{-}field}$*(M-P)*, at current intensities, 0.5-2.0 mA. E-field induced by 0.5mA tDCS in the deep M1 VOI shows linear correlation with the perfusion activity, $R^2$= 0.571 $p$=0.018 *(E)*.

Supplementary figure 5:

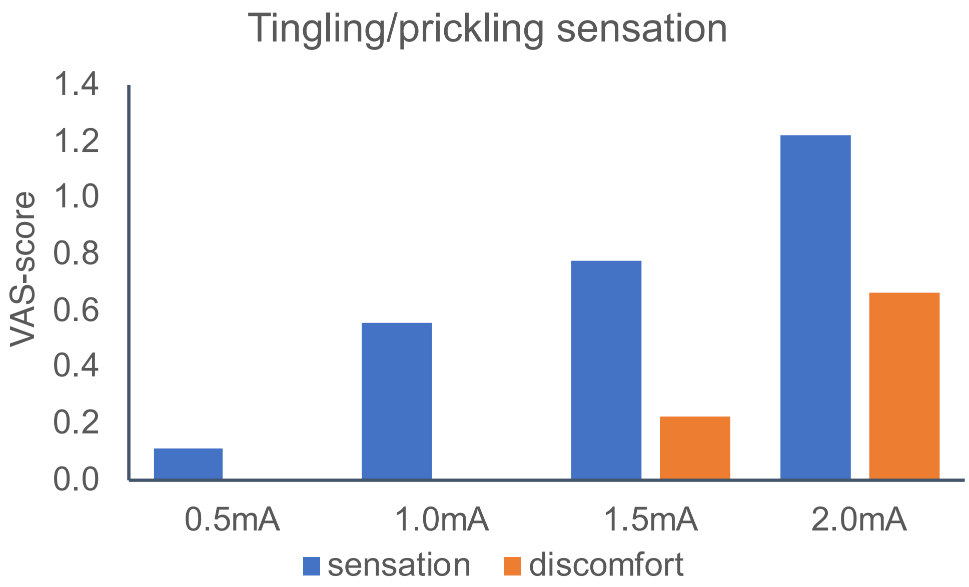


| | | | | | | | | | |
|---|---|---|---|---|---|---|---|---|---|
| | Did you experience a tingling/prickling sensation? | Not at all | 0 | 1 | 2 | 3 | 4 | 5 | Very much |
| | How uncomfortable was it? | Not at all | 0 | 1 | 2 | 3 | 4 | 5 | Very much |

**Supplementary figure 5:** Rating of somatosensory sensation during TDCS. Bar plot show average VAS-score rating of “tingling or pickling sensation” during all TDCS conditions (blue), and the corresponding level of discomfort (orange). VAS-score was rated on a 6-level scale ranging from “not at all” to “yes, very much”.

Supplementary table 1: VAS-score questionnaire:

Condition no. /tDCS intensity ___________

| | | | | |
|---|---|---|---|---|
| **Fatigue** | How fatigued are you right now? | Not at all | ☐☐☐☐☐☐ | Very much |
| **Malaise** | Are you uncomfortable right now? | Not at all | ☐☐☐☐☐☐ | Very much |
| **Nausea** | Did you feel nauseous? | No ☐ | | Yes ☐ |
| **Sensory** | Did you experience a tingling/prickling sensation? | Not at all | ☐☐☐☐☐☐ | Very much |
| | How uncomfortable was it? | Not at all | ☐☐☐☐☐☐ | Very much |
| **Phosphenes** | Did you see flickering lights? | Not at all | ☐☐☐☐☐☐ | Very much |
| | How uncomfortable was it? | Not at all | ☐☐☐☐☐☐ | Very much |
| **Metal taste** | Did you experience a metallic taste? | Not at all | ☐☐☐☐☐☐ | Very much |

**Supplementary table 1**: Questionnaire was repeated after each block of TDCS, a total of four times.
